# Characterization of alveolar epithelial lineage heterogeneity during the late pseudoglandular stage of mouse lung development

**DOI:** 10.1101/2022.01.05.475053

**Authors:** Matthew R. Jones, Lei Chong, Arun Reddy Limgapally, Jochen Wilhem, Meshal Ansari, Herbert B. Schiller, Gianni Carraro, Saverio Bellusci

## Abstract

The specification, characterization, and fate of alveolar type 1 and type 2 (AT1 and 2) progenitors during embryonic lung development remains mostly elusive. In this paper, we build upon our previously published work on the regulation of airway epithelial progenitors by fibroblast growth factor receptor 2b (Fgfr2b) signalling during early (E12.5) and mid (E14.5) pseudoglandular lung development. Here, we looked at the regulation by Fgfr2b signalling on alveolar progenitors during late pseudoglandular/early canalicular (E14.5-E16.5) development. Using a dominant negative mouse model to conditionally inhibit Fgfr2b ligands at E16.5, we used gene array analyses to characterize a set of potential direct targets of Fgfr2b signalling. By mining published single-cell RNA sequence (scRNAseq) datasets, we showed that these Fgfr2b signature genes narrow on a discreet subset of AT2 cells at E17.5 and in adult lungs. Furthermore, we demonstrated that Fgfr2b signalling is lost in AT2 cells in their transition to AT1 cells during repair after injury.

We also used CreERT2-based mouse models to conditionally knock-out the Fgfr2b gene in AT2 and in AT1 progenitors, as well as lineage label these cells. We found, using immunofluorescence, that in wildtype controls AT1 progenitors labeled at E14.5-E15.5 contribute a significant proportion to AT2 cells at E18.5; while AT2 progenitors labeled at the same time contribute significantly to the AT1 lineage. We show, using immunofluorescence and FACS-based analysis, that knocking out of *Fgfr2b* at E14.5-E15.5 in AT2 progenitors leads to an increase in lineage-labeled AT1 cells at E18.5; while the reverse is true in AT1 progenitors. Furthermore, we demonstrate that increased Fgfr signalling in AT2 progenitors reduces their contribution to the AT1 pool.

Taken together, our results suggest that a significant proportion of AT2 and AT1 progenitors are cross-lineage committed during late pseudoglandular development, and that lineage commitment is regulated in part by Fgfr2b signalling. We have characterized a set of direct Fgfr2b targets at E16.5 which are likely involved in alveolar lineage formation. These signature genes concentrate on a subpopulation of AT2 cells later in development, and are downregulated in AT2 cells transitioning to the AT1 lineage during repair after injury in adults.

Our findings highlight the extensive heterogeneity of alveolar cells by elucidating the role of Fgfr2b signalling in these cells during early alveolar lineage formation, as well as during repair after injury.

## Introduction

Around embryonic day (E) 9.5 in the mouse, a specialized region of *Nkx2-1* expressing endoderm of the anterior foregut evaginates, forming the anlage of the trachea and lung. During the ensuing embryonic and pseudoglandular stages of development (E9.5-E16.5), the growing multipotent epithelium undergoes branching morphogenesis, concomitant with the earliest differentiation and cell-fate specification of the various epithelial cell types. The developing epithelium quickly resembles a highly branched tree of proximal conducting airways, demarcated by SOX2 expression, and distal buds positive for SOX9 and ID2 (Rawlins et al., 2009). By the end of pseudoglandular development, and leading into the canalicular (E16.5-E17.5) and saccular stages (E17.5-postnatal (PN) day 4), an abundance of distal progenitors expressing markers for alveolar epithelial type 1 (AT1) and type 2 (AT2) cell lineages populates the nascent alveoli. These two epithelial lineages are essential for proper lung function: AT1 cell bodies form thin elongated extensions lining the apical sides of alveoli, providing a gas exchange interface between the alveolar lumen and the capillary plexus located on the basal epithelial surface; AT2 cells are cuboidal, are distributed around the perimeters of alveoli, and produce and secrete surfactant, which reduces epithelial surface tension thus preventing alveolar collapse. AT2 cells are also stem cells, capable of self-renewal or differentiation to an AT1 cell upon repair after injury (Barkauskas et al., 2013).

There is still much to elucidate concerning the journey of *Nkx2-1* positive multipotent progenitors towards an alveolar lineage specification and eventual commitment to an AT1 or AT2 fate. Most likely, a complex interplay of signalling pathways, biomechanical, and physical factors regulates the spatial and temporal emergence of alveolar progenitor cells during development (Jones et al., 2021; Li et al., 2018; Tang et al., 2018). At a basic level, however, it is still unclear when the alveolar lineage emerges, and what are the characteristics of the progenitor cell(s). This uncertainty is captured by two different, yet not necessarily exclusive, models of alveolar lineage specification: the bipotent progenitor model, and the early lineage specification model.

In the first, earlier model, it was proposed that oligopotent alveolar progenitors give rise to a progenitor population baptized “bipotent progenitor cells (BP)”. It was argued that these bipotent epithelial progenitor cells, which are detected at E16.5 at the end of the branching program, can self-renew or differentiate into either the AT1 (expressing the canonical marker *Pdpn*) or AT2 lineage (expressing the canonical markers *Sftpc* and *Sftpb*). Based on single-cell transcriptomic studies of the epithelium at different stages during lung development, a gene signature characteristic to each differentiated cell type was identified. The bipotent progenitor cells exhibit both AT1 and AT2 signatures, and, during differentiaiton, progressivley lose one or the other signature to become a mature alveolar cell (Treutlein et al., 2014).

An important limitation of the model placing BP cells as the hub for both AT1 and AT2 formation was that lineage tracing approaches for the BP cells were missing. In the second and more recent model which combined single cell transcriptomics along with lineage tracing, it was proposed that most mature AT1 and AT2 cells arise from unipotent progenitors, largely bypassing the BP cell type (Frank et al., 2019). Furthermore, it was suggested that alveolar progenitors form much earlier during development than anticipated, as early as E13.5. It was also shown using a dual transgenic approach that labeled BP did not significantly contribute to the alveolar lineage. One of the limitations of this study was the time points chosen did not allow to exclude a transient contribution of the committed AT2 progenitors to the AT1 lineage and vice-versa.

Continuing research on the mechanisms regulating alveolar lineage specification and differentiation will help refine our current models of this fundamental developmental process. For example, previous work from our lab has demonstrated a critical role for fibroblast growth factor (Fgf) 10 signalling via fibroblast growth factor receptor (Fgfr) 2b on AT2 lineage formation. For instance, when analyzed at PN3, heterozygous *Fgf10^+/-^* mice have a decreased ratio of AT2 cells over total EpCam-positive epithelial cells compared to wildtype controls (Chao et al., 2017), suggesting that Fgf10 signalling is required for AT2 lineage formation. Interestingly, after hyperoxia exposure, this AT2 pool gets normalized arising the interesting possibility that compensatory populations are getting engaged (Chao et al., 2017), Such compensatory mechanisms may involve the newly discovered sftpc-lineage labelled “injury activated alveolar progenitors (IAAPs) (Ahmadvand et al., 2021).

Prior to the initiation of the alveolar differentiation program, Fgfr2b signalling also tightly regulates branching morphogenesis (Jones et al., 2021). Research continues on the regulatory mechanisms downstream of Fgfr2b signalling which control, at the level of the multipotent epithelium, the transition from a branching program during the pseudoglandular stage (E11.5 to E16.5) to an alveolar program at the canalicular/saccular stages (E16.6 to E18.5). For example, it was shown that the forced activation of Fgf-dependent Kras expression in the distal epithelium suppressed the alveolar differentiation program by promoting, *via* SOX9, the branching program (Chang et al., 2013). This supports loss-of-function experiments which have demonstrated that SOX9 acts downstream of Fgf signaling to promote epithelial branching while preventing alveolar differentiation (Volckaert et al., 2013). It was also reported that continuous overexpression of *Fgf10* throughout lung development (from E11.5 to E18.5) prevents the formation of SOX2-positive bronchiolar progenitor cells while maintaining a high number of SOX9-positive cells (Volckaert et al., 2013). In addition, analysis of FGF10 overexpressors compared to control lungs indicated a drastic increased number of SFTPC-positive cells, suggesting that FGF10 could enhance the differentiation of the alveolar progenitor cells towards the AT2 lineage. FGF10 overexpression at later stages (from E15.5 to E18.5) also prevented the differentiation of already committed SOX2*-*positive bronchiolar progenitor cells to the ciliated-cell lineage, but facilitated instead their commitment to the P63-positive basal-cell lineage. Interestingly, in these experimental conditions, both SFTPC and PDPN-positive cells were observed, but the lack of quantification prevented determining whether the overexpression of FGF10 led to changes in the number of AT1 vs. AT2 cells (Volckaert et al., 2013). Loss of epithelial *Fgfr2* expression in alveolar progenitor cells using the *Id2-CreERT* driver line from E15.5 led to a decreased AT2/AT1 cell ratio at E18.5. A similar result was observed with pregnant females treated with PD0325901 (an extracellular-signal regulated kinase (ERK) inhibitor) at E15.5 and analyzed at E18.5 for the differentiation of ID2-lineage traced cells (Li et al., 2018). Furthermore, deletion of *Etv5*, a well-known Fgf target gene, in adult AT2 cells leads to the loss of AT2 markers and collateral acquisition of an AT1 signature in these cells (Zhang et al., 2017). Altogether, these results suggest that Fgf plays a critical role both in the differentiation of the alveolar progenitors towards the AT2 fate during development as well as in the maintenance of the AT2 cell signature.

More recently, we have published direct transcriptional targets of Fgfr2b signalling on distal epithelial cells during early (E12.5) and mid- (E14.5) pseudoglandular development (Jones et al., 2019, 2020). In this work, we had used a dominant negative transgenic mouse model to conditionally inhibit Fgfr2b signalling (*Rosa26^rtTA/rtTA^; Tg(tet(o)sFgfr2b)/+*) and to identify and characterize sets of target genes. These ‘Fgfr2b gene signatures’ correspond to the observed biological activites controlled by Fgfr2b. During early pseudoglandular development, Fgf10 acts directly on distal tip progenitors to control cell motility, adhesion and arrangement (Jones et al., 2019). This morphogenic role persists through mid-pseudoglandular development (E14.5), along with a shifting contribution to the increased proliferation of distal cells seen at this stage (Jones et al., 2020). A critical finding from these analyses was that significant impacts on AT2 differentiation markers were also observed. For example, the loss of Fgfr2b signalling led to a significant decrease in AT2 signature gene expression as early as E12.5 (Jones et al., 2019). This finding suggests that *bona fide* alveolar progenitors may exist earlier than currently appreciated, and that their regulation is concomitant with the dominant branching program.

In the current paper, we continue the effort to understand the targets downstream of Fgfr2b which control alveolar differentiation. Building upon our work at E12.5 and at E14.5 by looking here at late pseudoglandular/early canalicular (E16.5) development, we took a similar approach by inhibiting Fgfr2b ligand activity for nine hours using the dominant negative mouse model. We performed a gene array on mutant and littermate control lungs, and identified a set of downregulated genes. We validated, using published single-cell RNA-sequencing datasets of E17.5 and adult lung samples, the cell-types expressing these Fgfr2b signature genes. We also used Cre-based transgenic mouse models to conditionally inactivate Fgfr2b signalling in AT1 or AT2 progenitors, and to label those cells with a tomato reporter. With these models, we were able to lineage label putative alveolar progenitors just prior to E16.5 and analyse their commitment to both AT2 and AT1 lineages in control and loss of Fgfr2b signalling conditions. We also FACS-isolated E18.5 AT1 and AT2 cells derived from labeled AT2 progenitors at E14.5-E15.5, and conducted a gene array on these samples. We compared the results to our data derived from our global inhibition experiments. These findings help to inform and resolve current general models for alveolar lineage specification. In particular, we show that SFTPC-positive progenitors not only give rise to mature AT2 cells, but also to mature AT1, and that this contribution to the AT1 lineage is not minor. Our findings also help build a more particular model for the specific role of Fgfr2b signalling on alveolar progenitors during pseudoglandular lung development.

## Results

### Global inhibition of Fgfr2b ligands reveals a set of potential direct targets of Fgfr2b signalling at E16.5

To assess the global transcriptional impacts of Fgfr2b signalling in late pseudoglandular / early canalicular (E16.5) embryonic mouse lungs, we utilized a previously described and validated transgenic mouse model: *Rosa26^rtTA/rtTA^*; *Tg(Tet(o)sFgfr2b)/+* (Danopoulos et al., 2013; Jones et al., 2019, 2020; Taghizadeh et al., 2020). In this system, a soluble form of the Fgfr2b receptor controlled by a Tet(o) promoter (*Tg(Tet(o)sFgfr2b)*) is transcribed upon activation by doxycycline of the ubiquitously expressed rtTA (*Rosa26^rtTA^*). Here, we injected E16.5 timed-pregnant females carrying experimental (*Rosa26^rtTA/rtTA^*; *Tg(Tet(o)sFgfr2b)/+*) and littermate controls (*Rosa26^rtTA/rtTA^*; *+/+*) with doxycycline intraperitoneally (Dox-IP) and harvested the embryos and their lungs nine hours later (Fig. 1A).

**Figure 1:**
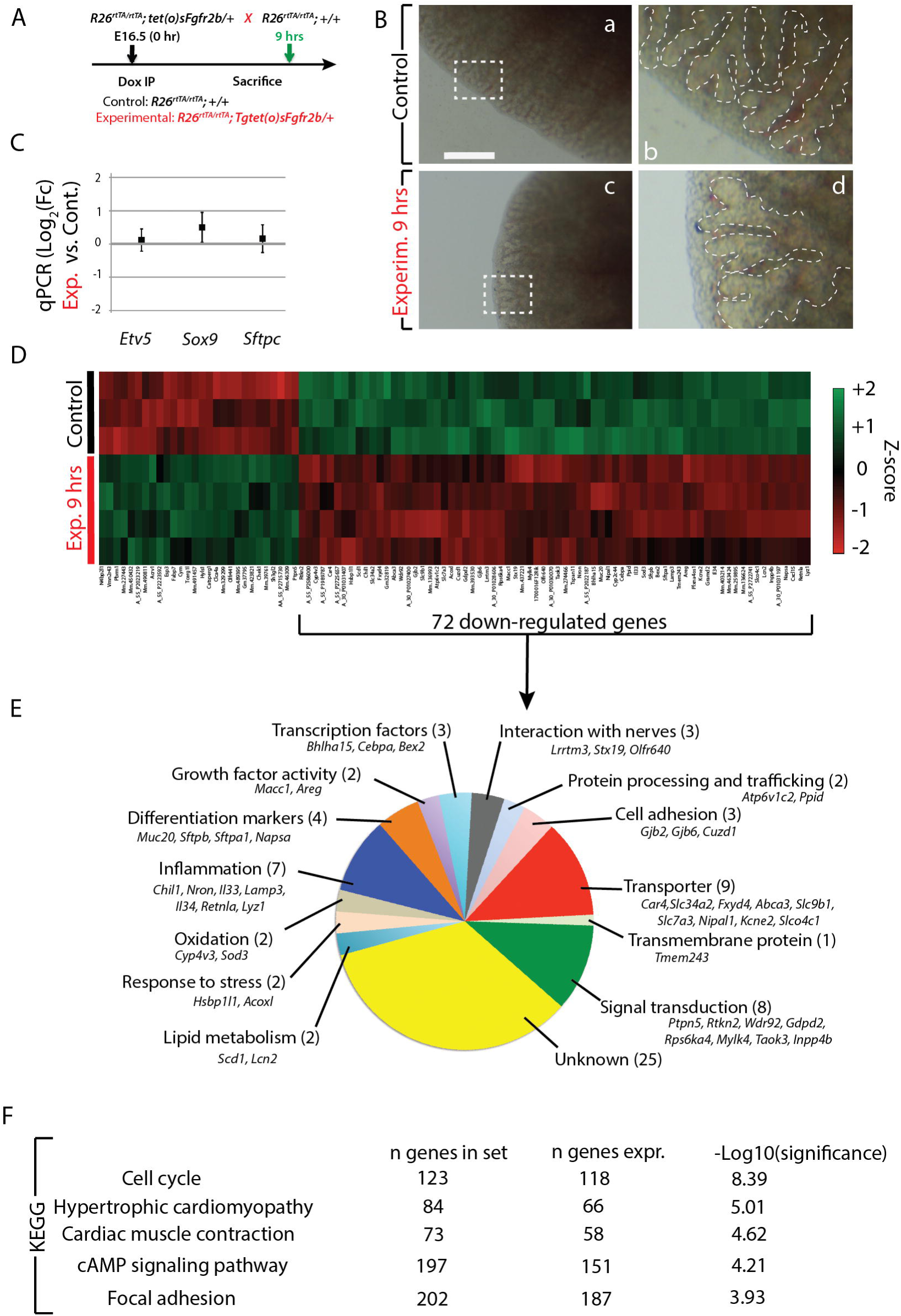
Transcriptomic changes upon Fgfr2b signalling inhibition at E16.5 + 9 hours. **(A)** Experimental design. E16.5 littermate experimental and control lungs were collected 9 hours after a single Dox-IP injection. **(B)** Gross lung morphology appears similar between control (a and b) and experimental (c and d) lungs. **(C)** Gene expression analysis by qPCR of differentiation markers *Etv5*, *Sox9*, and *Sftpc*. **(D)** Heatmap of the top 100 genes (according to p-value) differentially expressed after 9 hours Fgfr2b inhibition. 72 of those genes are down-regulated, representing potential direct targets of Fgfr2b signalling. **(E)** Gene ontology of the 72 down-regulated genes found in ‘D’. **(F)** Top five KEGG pathways regulated from the complete gene array.

Macroscopically, the phenotypes of control and experimental lungs showed little difference (Fig. 1B), nor was there an appreciable effect on the expression of the epithelial differentiation markers *Etv5*, *Sox9*, and *Sftpc* as determined by qPCR (Fig. 1C). A gene array was conducted on whole lung homogenates from control (n=3) and experimental (n=4) embryos. A heatmap of the top 100 genes (according to p-value) differentially regulated between experimental and control samples is shown in Fig. 1D. From these 100 genes, 72 were down-regulated in experimental versus control lungs; these genes constitute the Fgfr2b signature at E16.5 (see Fig. S1). Gene ontology (Fig. 1E) and KEGG pathway analyses (Fig. 1F) of these 72 genes indicate that a number of biological processes are regulated by Fgfr2b signalling at this time point. These include, in order from high to low of the number of genes involved, cellular transporting (9 genes), signal transduction (8), inflammation (7), markers of differentiation (4), transcriptional regulation (3), cell adhesion (3), interaction with nerves (3), protein processing and trafficking (2), lipid metabolism (2), response to stress (2), oxidation (2), growth factor activity (2), and transmembrane proteins (1). The pathways primarily affected by these genes include cell cycle regulation, cAMP signaling, focal adhesion, and, interestingly, hypertrophic cardiomyopathy and cardiac muscle contraction. In the context of alveolar lineage formation, these latter two pathways are likely linked to the migration of AT2 progenitors out of the lumen to escape the physical forces applied by the amniotic fluid, a process regulated by Fgfr2b-mediated actin/myosin-dependent mechanisms (Li et al., 2018).

### AT1 and AT2 progenitor lineage labeling suggests a level of cross-lineage contribution

It has been argued that AT1 and AT2 progenitor cell populations are largely unipotent as early as E13.5 during mouse lung development (Frank et al., 2019). This contrasts with the earlier model proposing that the majority of mature AT cells pass through a bipotential progenitor well after E13.5 (Treutlein et al., 2014). To address this issue, we investigated the lineage-commitment of early alveolar progenitors using our mouse models. We labeled AT1 and AT2 progenitor cells during the late pseudoglandular stage using *Hopx^CreERT2/+^; tdTomato^flox/flox^* (n=3) and *Sftpc^CreERT2/+^*; *tdTomato^flox/flox^* (n=3) transgenic mouse lines, respectively. We labeled cells via two tamoxifen IP (Tam-IP) injections (E14.5 and E15.5) and assessed the contribution of lineage labeled cells to each AT population at E18.5 (Fig. 2A). Assessing the expression of SFTPC and of HOPX proteins via immunofluorescence staining in the *Sftpc^CreERT2^* line, it was observed that around 75% and 20% of labeled (RFP^Pos^) cells were SFTPC^Pos^ and HOPX^Pos^, respectively (Fig. 2B and C). The expression of labeled SFTPC^Pos^ and of HOPX^Pos^ cells in the *Hopx^CreERT2^* line was essentially the reverse of what was found in the *Sftpc^CreERT2^* line: around 70% and 20% of RFP^Pos^ cells were HOPX^Pos^ and SFTPC^Pos^, respectively (Fig. 2D and E). These data demonstrate that a significant proportion of mature AT cells (around 20%) derive from inter-lineage progenitors.

**Figure 2:**
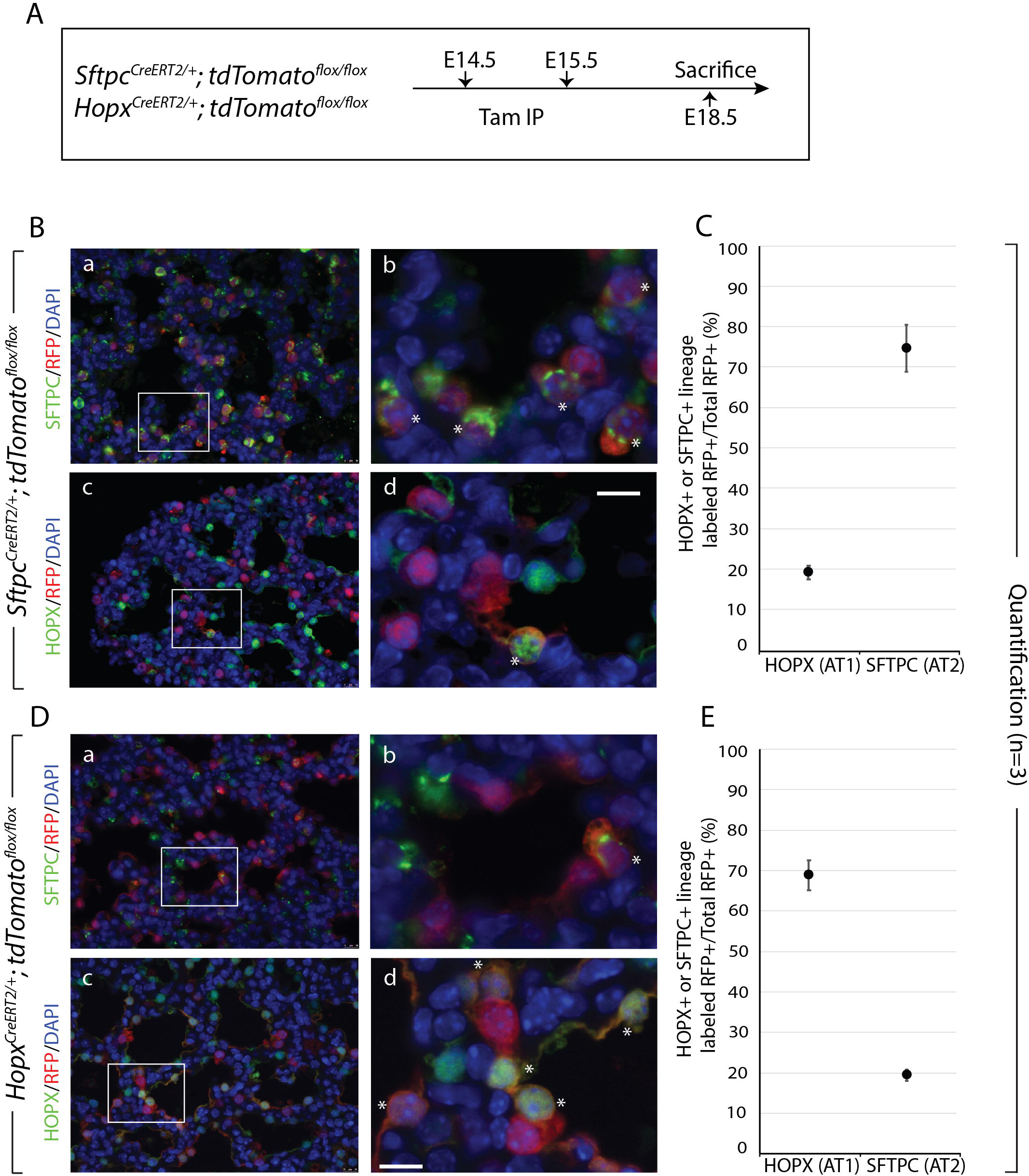
Lineage tracing of AT2 (*Sftpc*) and AT1 (*Hopx*) progenitors suggests inter-lineage contributions. **(A)** Transgenic mice carrying a *CreERT2* allele downstream of a *Sftpc* promoter or a *Hopx* promoter were used to label AT2 or AT1 progenitors, respectively, by recombining a floxed *TdTomato* reporter. Timed-pregnant females were injected interperitoneally with tamoxifen (Tam-IP) at E14.5 and E15.5, and embryonic lungs were harvested just prior to birth at E18.5. **(B)** Lineage labeled *Sftpc^CreERT2/+^*; *tdTomato^flox/flox^* embryonic lungs were immunostained for SFTPC (a and b) and HOPX (c and d) to assess the contribution of labeled cells to the AT2 and AT1 lineages, respectively (see asterisks in panels b and d for examples of lineage labeled AT2 and AT1 cells, respectively). **(C)** Labeled AT1 and AT2 cells were counted from multiple images taken from three independent samples (n=3) and are displayed as a percentage of the total number of Tomato RFP^pos^ cells. Labeled HOPX (AT1) cells compose around 20% of the total RFP^pos^ population, whereas labeled (SFTPC) AT2 cells compose around 75%. **(D)** Lineage labeled *Hopx^CreERT2/+^*; *tdTomato^flox/flox^* embryonic lungs were immunostained for SFTPC (a and b) and HOPX (c and d) to assess the contribution of labeled cells to the AT2 and AT1 lineages, respectively (see asterisks in panels b and d for examples of lineage labeled AT2 and AT1 cells, respectively). **(E)** Labeled HOPX (AT1) cells compose around 70% of the total RFP^pos^ pool, whereas labeled (SFTPC) AT2 cells compose around 20%.

### Cell autonomous deletion of *Fgfr2b* in AT2 progenitors pushes them toward the AT1 lineage

To investigate the role of Fgfr2b signalling specifically on AT2 progenitor cell fate, we utilized a *Sftpc^CreERT2/+^*; *tdTomato^flox/flox^*; *Fgfr2b^flox/flox^* line. This line labels AT2 (*Sftpc*) progenitors while simultaneously excising the IIIb isoform of the *Fgfr2* gene (De Moerlooze et al., 2000).

In our first experiment, we Tam-IP injected at E14.5 and E15.5 timed-pregnant females carrying either control (*Sftpc^CreERT2/+^*; *tdTomato^flox/flox^*) (n=7) or experimental (*Sftpc^CreERT2/+^*; *tdTomato^flox/flox^*; *Fgfr2b^flox/flox^*) (n=2) embryos and harvested the lungs at E18.5 (Fig. 3A). We then conducted FACS-based isolation of RFP-labeled AT1 (EpCAM^Pos^, RFP^Pos^, T1α^Pos^) and AT2 (EpCAM^Pos^, RFP^Pos^, T1α^Neg^) cells from these samples (see Fig. S2A). As Figure 3B shows, the percentage of labeled AT2 cells relative to the RFP^Pos^ epithelium (EpCAM^Pos^, RFP^Pos^) dropped from around 80% in controls to 70% in experimental lungs, while the percentage of labeled AT1 cells trended upwards from 15% to 20%.

**Figure 3:**
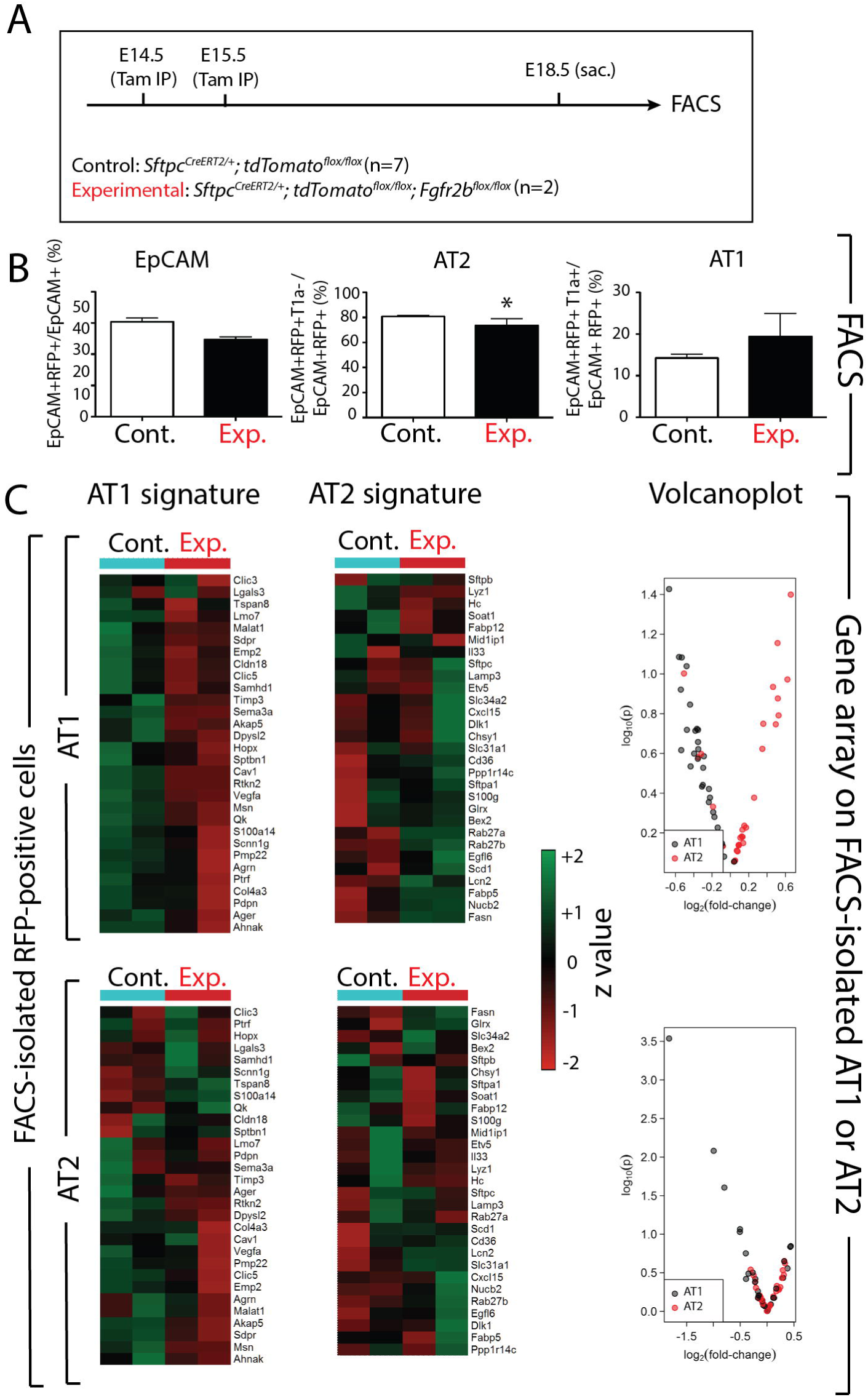
Cell autonomous deletion of *Fgfr2b* in AT2 progenitors results in decreased AT1 and increased AT2 signature expressions in isolated labeled AT1 cells. **(A)** Experimental design. Pregnant females carrying either control or experimental embryos were injected with TAM-IP at E14.5 and E15.5, and embryonic lungs were harvested at E18.5. **(B)** Control and experimental lungs were processed for FACS analysis and isolation of cells. Tomato RFP-labelled epithelial cells (RFP^pos^ EpCAM^pos^) were sorted, and AT1 and AT2 cells were isolated based on the expression of the AT1 marker T1α (Podoplanin). A slight decrease in the proportion of tomato-labelled epithelial cells is observed in experimental versus control lungs (EpCAM^pos^ RFP^pos^/EpCAM^pos^), along with a small yet significant decrease in labelled AT2 cells (EpCAM^pos^ RFP^pos^ T1α^neg^/EpCAM^pos^ RFP^pos^). On the other hand, an increasing trend is seen in the proportion of AT1 cells from experimental versus control lungs (EpCAM^pos^ RFP^pos^ T1α^pos^/EpCAM^pos^ RFP^pos^). **(C)** Gene arrays on the isolated RFP^pos^ AT1 and AT2 cells were conducted, and heatmaps, as well as corresponding volcano plots, showing the relative expression of the AT1 and AT2 gene signatures in each cell population are presented. Of significance is the global downregulation and upregulation of AT1 and AT2 signature genes, respectively, in isolated AT1 cells.

We subsequently conducted a gene array on the FACS-isolated AT1 and AT2 cells from each condition (control and experimental) (Fig. S2B-D). As a quality control measure, we compared AT1 and AT2 gene signature expressions (as published by Treutlein et al. (2014)) in the isolated AT1 cells against the signature expressions in isolated AT2 cells, in each condition (Fig. S2E). As expected, the isolated AT1 cells showed an increased AT1 signature expression compared to the isolated AT2 cells, while the AT2 cells displayed an increased AT2 signature expression in control and experimental (Fig. S2E) lungs confirming that our FACS-based approach accurately enriched AT1 and AT2 cells

Finally, we compared AT1 and AT2 gene signature expressions in isolated AT1 and AT2 cells from control versus experimental lungs (Fig. 3C). Interestingly, in the isolated AT1 cells, the AT1 signature showed a lower expression in experimental compared to control lungs, while the AT2 signature actually increased in experimental AT1 cells compared to controls. In the isolated AT2 cells, a slight decrease was observed in the AT1 signature between control and experimental lungs, while little change was observed in the AT2 signature expression. Taken together, these data suggest that AT2 progenitors which lose Fgfr2b signalling activity might escape from the AT2 lineage and move into the AT1 lineage (as evidenced by the increase in labeled AT1 cells as well as the higher AT2 signature seen in these cells in experimental conditions).

To further investigate these surprising results from the FACS data, we assessed the lungs by qPCR and immunofluorescence staining at E18.5 (Fig. 4A). Figure 4B shows immunofluorescence antibody staining for HOPX (a-d) as well as SFTPC (e-h), in control (n=3) and experimental (n=3) lungs. Asterisks indicate lineage-labeled AT1 or AT2 cells. These samples were manually quantified (Fig. 4C). The percentage of double positive AT1 cells (HOPX^Pos^ RFP^Pos^) over the total pool of lineage labeled cells (RFP^Pos^) increased from around 20% to 28% in experimental lungs (Fig. 4Ca), while the level of labeled AT2 cells (SFTPC^Pos^ RFP^Pos^) remained around the same (78%) in both control and experimental conditions (Fig. 4Cb). The percentage of lineage labeled AT1 or AT2 cells relative to the pool of total AT1 or AT2 cells decreased (Fig. 4Db and d), likely as a consequence of reduced proliferation of lineage labeled cells in the experimental condition.

**Figure 4:**
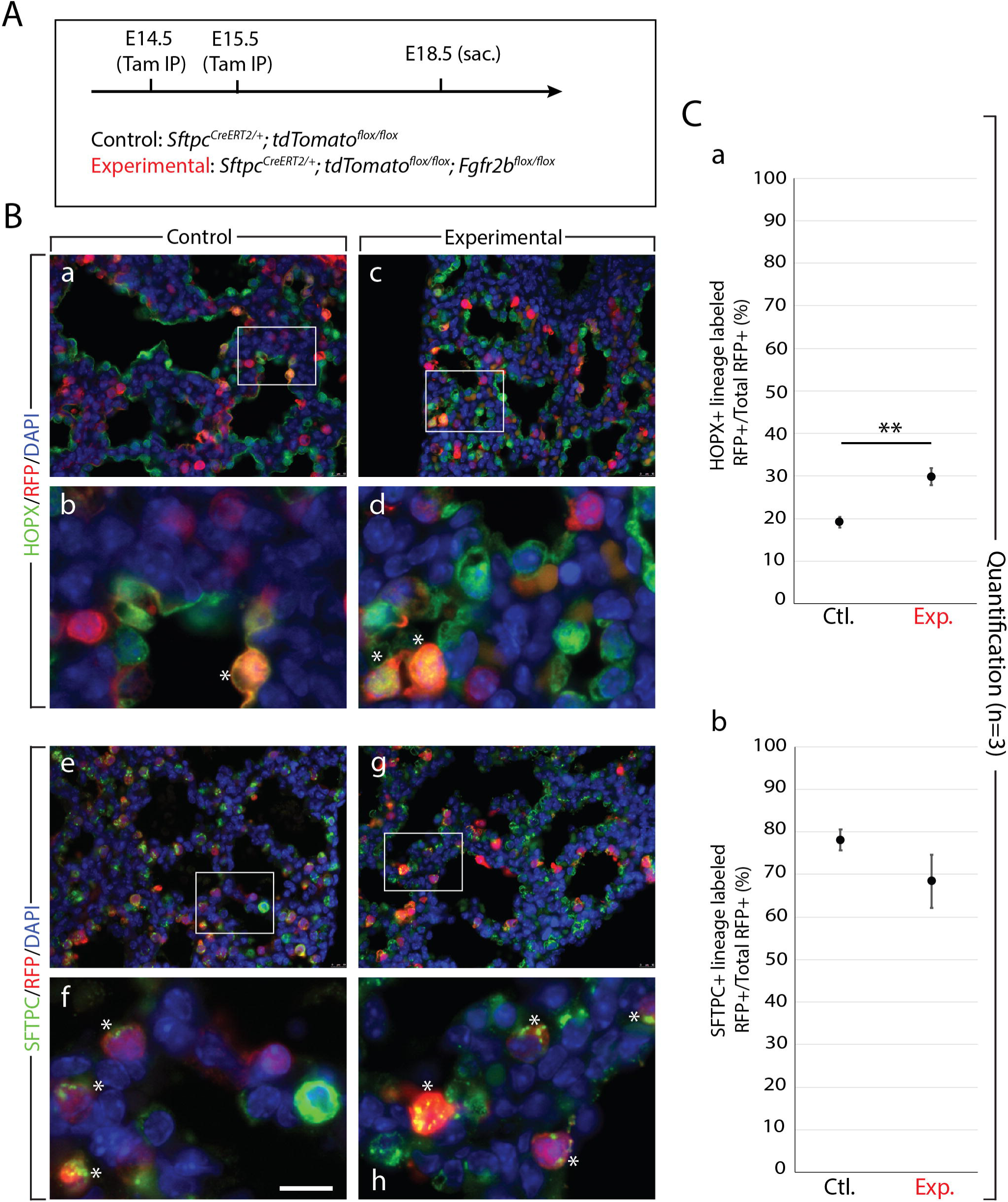
Analysis of Fgfr2b deletion in AT2 progenitors. **(A)** Experimental design. Pregnant females carrying either control or experimental embryos were injected with TAM-IP at E14.5 and E15.5, and embryonic lungs were harvested at E18.5. **(B) qPCR (C)** Immunofluorescence staining of either AT1 cells (HOPX; panels a-d) or AT2 cells (SFTPC; panels e-h) in control (a, b and e, f) and experimental (c, d and g, h) lineage-labeled (RFP) samples. Asterisks indicate double positive cells. *Scale bar:* 30 µm (a, c, e, g); 75 nm (b, d, f, h). **(D)** Quantification of samples from (C). Graphs show the percentage of double positive cells either from the pool of total lineage-labeled cells (a and c) or from the pool of AT1 or AT2 cells (b and d). Loss of Fgfr2b in AT2 progenitors results in an increase from around 20% to 28% in the percentage of labeled cells which are AT1 (HOPX^pos^, graph a), whereas the percentage of labeled cells which are AT2 remains around 78% (this will likely change with more samples analyzed) in both control and experimental lungs (graph c). Graphs b and d show that the percentage of double positive cells belonging to either lineage decreases in experimental lungs.

### Cell autonomous deletion of *Fgfr2b* in AT1 progenitors pushes them toward the AT2 lineage

Next, we investigated the role of Fgfr2b signalling specifically on AT1 progenitor cells by utilizing *Hopx^CreERT2/+^*; *tdTomato^flox/flox^*; *Fgfr2b^flox/flox^* mice to label and delete *Fgfr2b* expression in AT1 progenitors. Following Tam-IP at E14.5 and E15.5 we assessed control and experimental lungs by immunofluorescence staining at E18.5 (Fig. 5).

**Figure 5:**
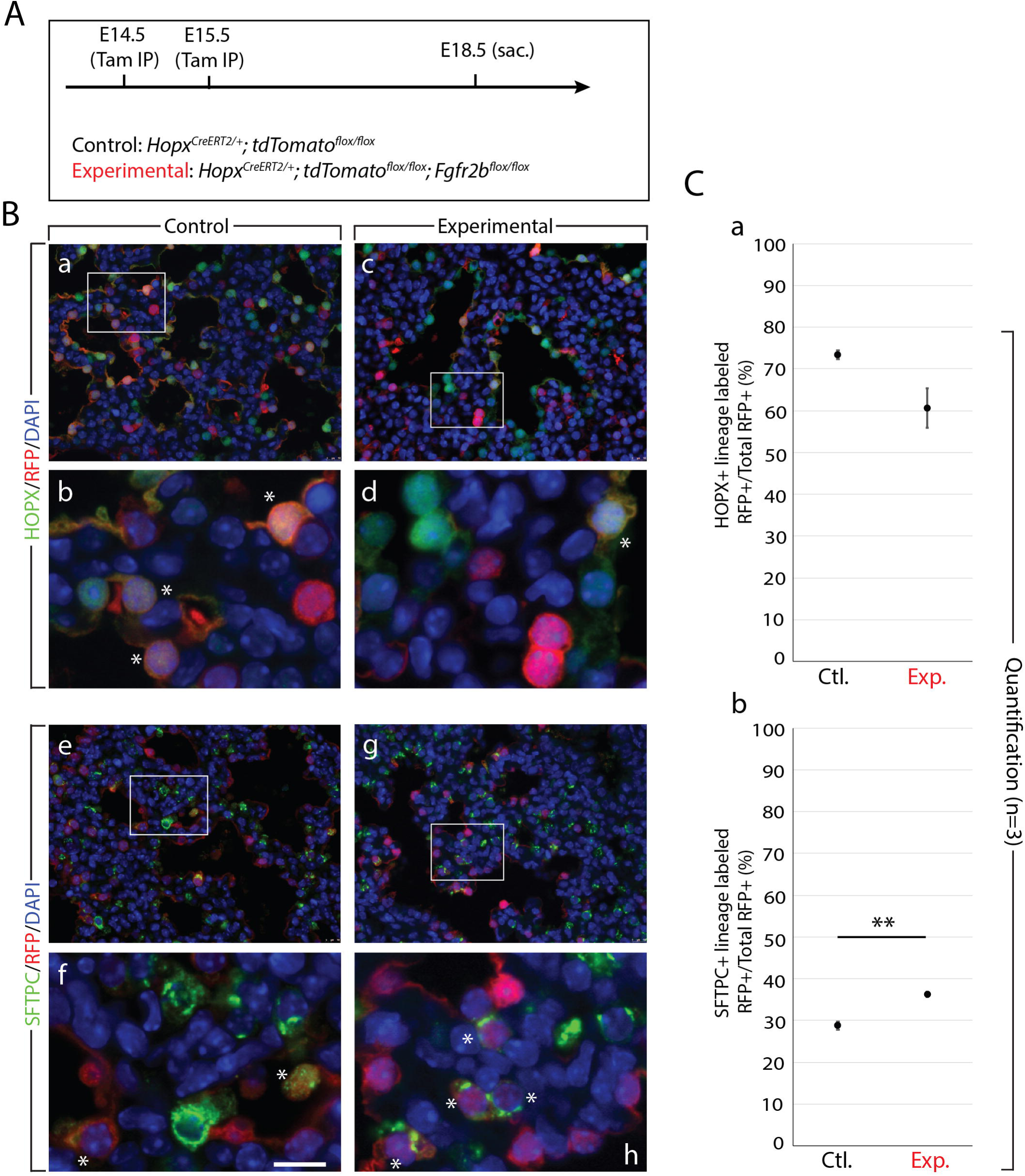
Analysis of Fgfr2b deletion in AT1 progenitors. **(A)** Experimental design. Pregnant females carrying either control or experimental embryos were injected with TAM-IP at E14.5 and E15.5, and embryonic lungs were harvested at E18.5. **(B)** qPCR-based gene expression analysis on whole lung homogenates shows that while *Fgfr2b* and *Sftpc* shows little regulation in experimental lungs, the expression of *Aqp5*, another canonical AT1 marker, is significantly reduced. **(C)** Immunofluorescence staining of either AT1 cells (HOPX; panels a-d) or AT2 cells (SFTPC; panels e-h) in control (a, b and e, f) and experimental (c, d and g, h) lineage-labeled (RFP) samples. Asterisks indicate double positive cells. *Scale bar:* 30 µm (a, c, e, g); 75 nm (b, d, f, h). **(D)** Quantification of samples from (C). Graphs show the percentage of double positive cells either from the pool of total lineage-labeled cells (a and c) or from the pool of total AT1 or AT2 cells (b and d). Loss of Fgfr2b in AT1 progenitors results in a decrease from around 73% to 61% in the percentage of labeled cells which are AT1 (HOPX^pos^, graph a), whereas the percentage of labeled cells which are AT2 increases with high statistical significance from 29% to 36% (graph c). Graphs b and d show that the percentage of double positive cells belonging to either lineage decreases in experimental lungs.

Figure 5B shows immunofluorescence antibody staining for HOPX (a-d) and SFTPC (e-h) in control and experimental lungs. Asterisks indicate lineage labeled AT1 or AT2 cells. Manual quantification of images is reported in Figure 5C. The percentage of labeled AT1 cells (HOPX^Pos^ RFP^Pos^) from the total RFP pool decreased in experimental lungs (Fig. 5Ca), whereas an increase in labeled AT2 cells over total labeled cells (SFTPC^Pos^ RFP^Pos^/RFP^Pos^) was observed (Fig. 5Cb). Consistent with the previous experiment (Fig. 4), lineage labeled cells as a percentage of total AT1 or AT2 cells are decreased in experimental samples, likely as a result of the overall decrease in proliferation of RFP^Pos^ cells (Fig. 5Db, d).

Taking the results from figures 3, 4, and 5 together, our data strongly suggests that labeled AT1 or AT2 progenitors preferentially switch lineages after cell autonomous loss of Fgfr2b signalling.

### Fgfr2b responsive genes during pseudoglandular development narrow on a subpopulation of AT2 cells at E17.5

Using the same transgenic mouse model as we use in this paper to globally inhibit Fgfr2b ligands (*Rosa26^rtTA/rtTA^*; *Tg(Tet(o)sFgfr2b)/+*), we have previously published Fgfr2b responsive genes during the early (E12.5) and mid (E14.5) pseudoglandular stages of lung development (Jones et al., 2019, 2020). Together with the 72 genes found in this paper to be downregulated upon Fgfr2b ligand inhibition (recall Fig. 1D), these shifting transcriptional targets (from E12.5 to E16.5) likely reflect the shifting biological roles played by Fgfr2b signalling at these different time points. A shortcoming of this global approach, however, is the inability to accurately characterize and assess the relative contribution to the global response from the different epithelial cell types. Which cell type responds most to Fgfr2b signalling? Are there important cell subpopulations to consider? How do responsive populations shift over developmental space and time?

One way to tackle this lack of information is to data-mine published single-cell RNA-sequencing (scRNA-Seq) datasets at different developmental time-points to obtain gene expression data for genes of interest. In this vein, we mined the recently published scRNA-Seq data from Frank et al. (2019), which contains transcriptome data from 7,106 individual Nkx2-1-positive epithelial cells obtained from E17.5 mouse lungs (GEO accession code GSE113320). Of the 11 categories denoted by the authors, four are of interest to us in this paper: AT1 precursor (preAT1), AT1, AT2 precursor (preAT2), and AT2 (Frank et al., 2019). These groups are depicted and colour-coded in Supplementary Figure S3A, along with the expressions of the AT2 markers *Sftpc* and *Lamp3,* and the AT1 markers *Ager* and *Aqp5* (Fig. S3B), as well as the expressions for *Fgfr2* and the well-established downstream effector of Fgfr2b signalling, *Etv5* (Fig. S3C). Note the comparable expressions of *Fgfr2* in preAT2 and AT2 cells, but also the presence of this gene in preAT1 cells. Finally, we assessed the expression in the four clusters of the signature gene sets we discovered at E12.5 (Jones et al., 2019), at E14.5 (Jones et al., 2020), and at E16.5 (this paper) (Fig. S3D-G). As the gene sets approach the time point of the sequenced single cells (E17.5), the expression profile becomes increasingly concentrated in a subset of the preAT2 and AT2 clusters (Fig. S3G). This finding suggests that the Fgfr2b signalling target genes discovered using our global Fgfr2b ligand inhibition model at E16.5 are predominantly expressed by a subset of the AT2 lineage at E17.5.

Following the identification of a subset of AT2 cells which highly express the Fgfr2b responding genes at E16.5, we further divided the AT2 cluster into two subclusters, termed cluster A and cluster B (Fig. 6A). We created a heatmap showing the top 50 differentially upregulated genes in each cluster (according to average Log_10_foldchange) (Fig. 6B). While canonical markers of AT2 cells, such as *Sftpc*, *Sftpb*, *Sftpa1*, and *Lamp3* are expressed in each subcluster, they are much more highly expressed in cluster B (Fig. 6C), suggesting that cluster B might represent a more mature pool of AT2 cells. Indeed, the genes most highly expressed in cluster A are predominately *markers of the AT1* lineage. Finally, while *Fgfr2* and *Etv5* expressions are scattered somewhat uniformly throughout clusters A and B (Fig. 6C), the targets of Fgfr2b signalling from E14.5 to E16.5 concentrate ever more narrowly in a subcluster of cluster B. The characterization of this Fgfr2b responding subcluster remains to be determined.

**Figure 6:**
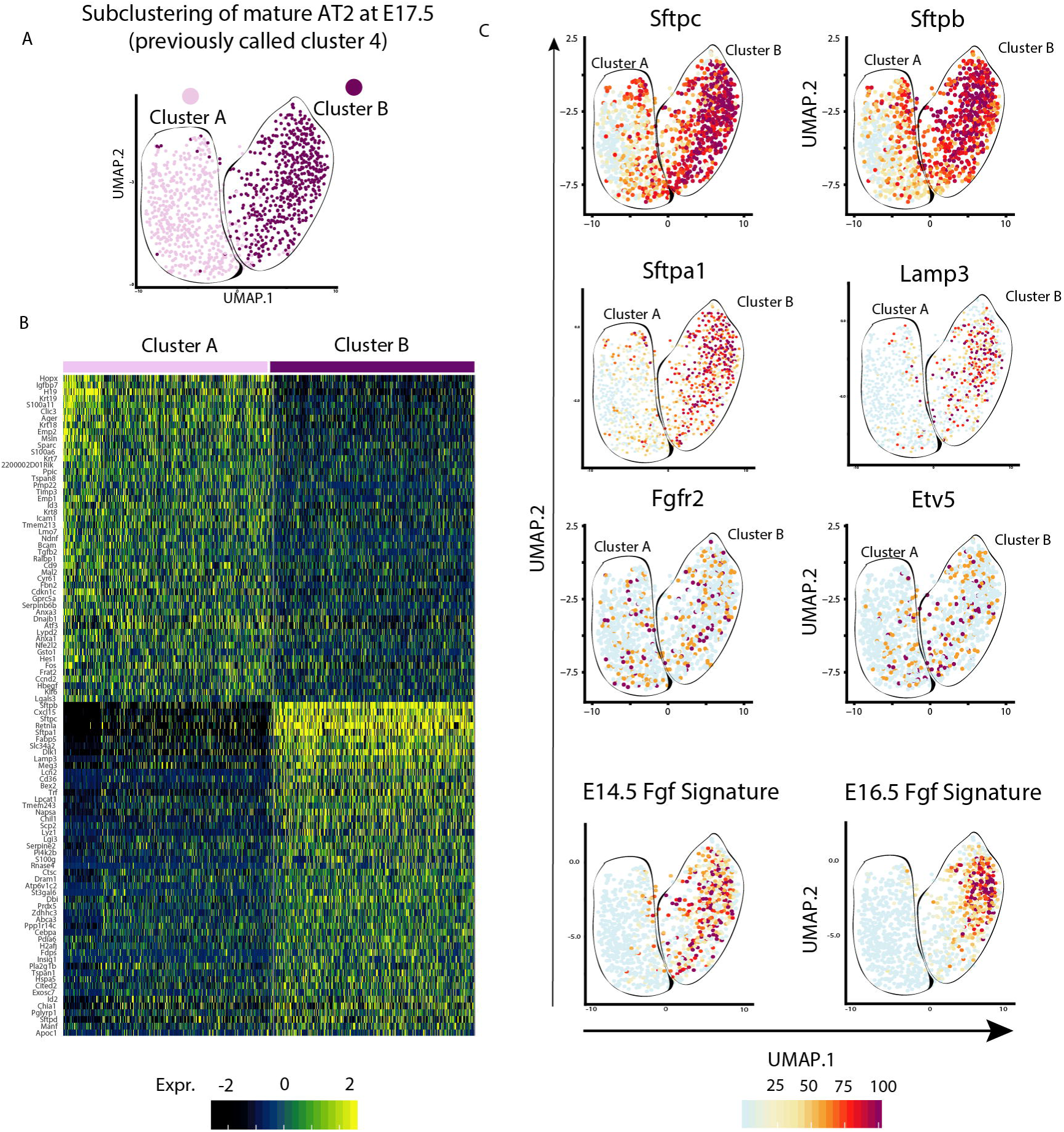
Fgfr2b signature concentrates into a subcluster of mature AT2 cells at E17.5. **(A)** Subclustering of mature AT2 cells as identified by scRNA-seq at E17.5 (Frank et al., 2019) (GEO GSE113320). The cells cluster into two discrete subclusters, termed A and B. **(B)** Heatmap showing the two 50 upregulated genes in each subcluster, according to Log_10_foldchange. While subcluster B is enriched in canonical AT2 marker genes, subcluster A shows enrichment in AT1 markers. **(C)** The expression profiles of select genes in subclusters A and B shows high expression of markers for mature AT2 cells in cluster B: *Sftpc, Sftpb, Sftpa1* and *Lamp3*. Note how *Fgfr2b* and *Etv5*, markers for Fgfr2b signalling, are scattered throughout subclusters A and B, while the Fgfr2b signalling signatures determined by our lab at E14.5 and at E16.5 concentrate on a subcluster of cluster B.

### AT2 cells lose Fgfr2b signalling as they transition to AT1 during repair after injury

A recent publication assessed the cell-type specific transcriptomic sequences involved in the regeneration of the airway after bleomycin-induced injury (Strunz et al., 2020). Using a combination of longitudinal scRNAseq data, RNA trajectory modeling in pseudo-time, as well as lineage tracing experiments, the authors demonstrated that during lung regeneration, populations of airway and AT2 stem cells converged on a transitional alveolar cell-type characterized by high *keratin 8* (*Krt8*) expression. These *Krt8*-positive airway differentiation intermediate (Krt8+ ADI) cells eventually gave rise to AT1 cells during repair of the gas exchange surface.

The scRNA-seq dataset published by Strunz et al. (2020) (GEO accession code GSE141259) provides useful cellular classifications, such as activated AT2 cells (characterized by injury-induced gene expression profiles), which are a subcluster of AT2s (marked by high *Sftpc* expression), as well as airway differentiation intermediate (ADI) cells. These subclusters are fully transcriptionally defined, and are placed in temporal context as transition states between mature AT2 and AT1 cells after injury. Since alveolar regeneration recapitulates earlier developmental stages, it is likely that these transitional states seen in adult lungs during repair after injury are also found in embryonic stages during normal development. Furthermore, as Fgfr2b signalling plays a critical role in the differentiation of alveolar cells during normal development, it is likely to also play a role during repair after injury. Therefore, we asked what is the level and where is the expression of the Fgfr2b signatures previously found at E12.5, at E14.5, and in this paper at E16.5 in adult lung cells following bleomycin injury using the scRNAseq dataset.

Figure 7A reproduces the UMAP from the original Strunz et al. (2020) study and highlights the Fgfr2b signatures we found at E12.5, at E14.5, and at E16.5. As can be seen, the signatures from each embryonic developmental stage increasingly concentrate on the AT2 and AT2-activated cell populations found during repair after bleomycin-induced injury (Fig. 7B and C). Note that the E12.5 signature is nearly equally present in AT1, AT2, and AT2-activated cells (left column in Fig.7B). As development progresses through E16.5, Fgfr2b signalling concentrates on the AT2 and AT2-activated populations (Fig. 7C), with a negligible scattering of expression in Krt8+ ADI cells and mature AT1 cells. Indeed, as AT2 stem cells transition through a Krt8+ ADI state towards an AT1 fate, Fgfr2b signature gene expression is significantly and drastically downregulated (Fig.7D). These findings suggest that developmentally-relevant Fgfr2b signalling is active in AT2 cells during homeostasis and that it is sustained in activated AT2 cells during repair after injury. Fgfr2b signaling might function to prevent AT2 cells from transitioning to an AT1 fate.

**Figure 7:**
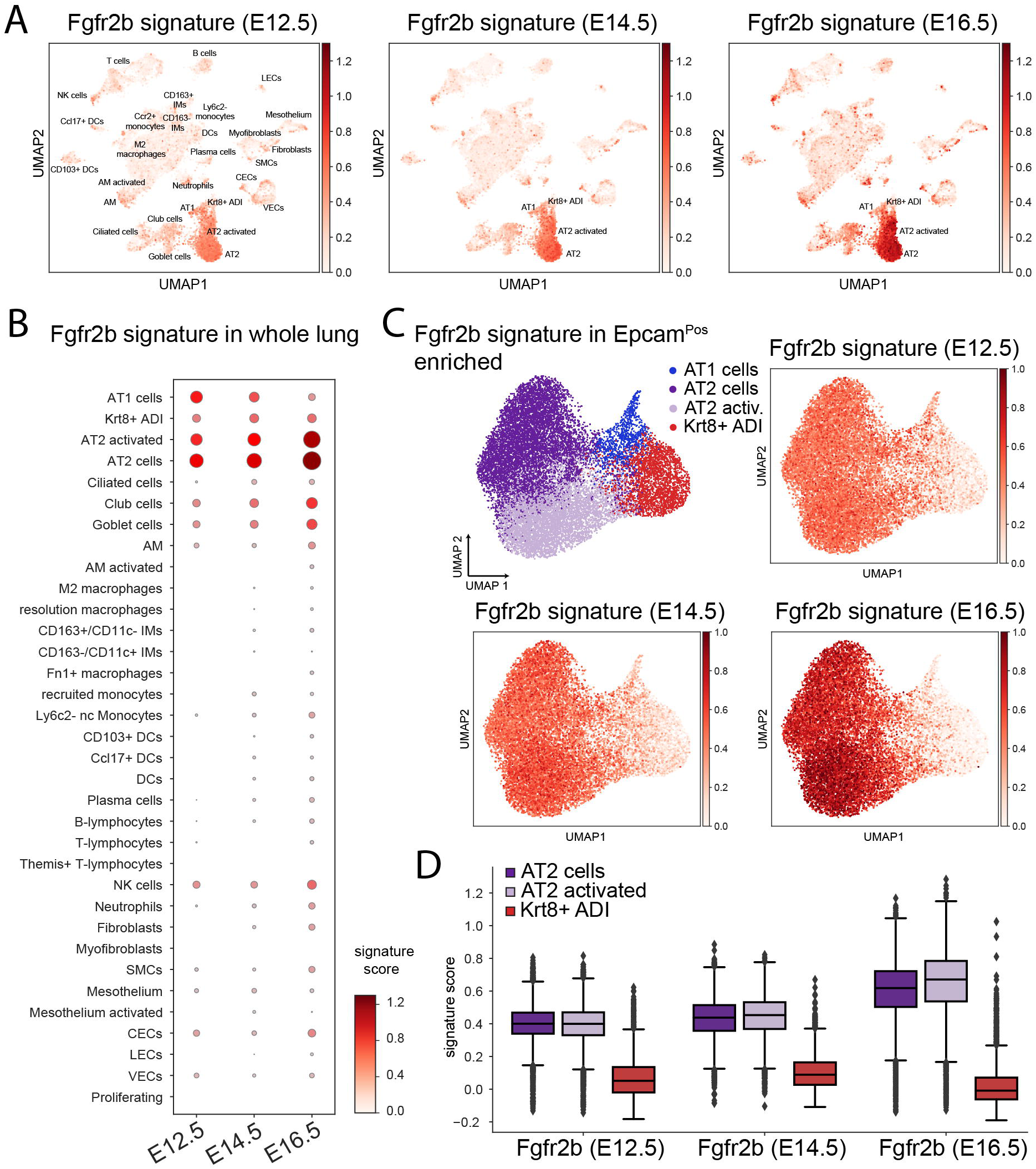
Fgfr2b signalling is lost in AT2 cell transition to AT1 during repair after injury. **(A)** scRNA-seq data from Strunz et al. (2020) (GEO accession code GSE141259) reanalyzed using the Fgfr2b gene signatures found by our lab at E12.5, E14.5, and E16.5. Note the increasing expression of these signature genes as embryonic development progresses in the AT2 clusters. **(B)** Dot plot showing the different cell categories from ‘A’ and the relative Fgfr2b signature scores at each embryonic timepoint in each category. **(C)** The Fgfr2b signature from different embryonic time points (E12.5, E14.5 and E16.5) concentrate in the AT2 clusters of Epcam^pos^ enriched alveolar cells. **(D)** Box and whisker plots of the AT2 clusters, along with the Krt8^pos^ ADI cluster found in ‘C’ showing the Fgfr2b signature score at the given time points. While each embryonic Fgfr2b signature remains relatively high in the AT2 cells, each is virtually non-existent in the transitional Krt8^pos^ ADI cluster.

## Discussion

### Cross contribution of AT1 and AT2 progenitors to the opposite lineages

The characterization and relative contribution of the early alveolar progenitors to the AT1 and AT2 lineages remains to be securely established. What proportion of mature alveolar epithelial cells pass through a bipotential progenitor, for example? Do bipotential cells represent a major progenitor pool, as was earlier argued (Treutlein et al., 2014), or a minor subsidiary contributor to mature alveolar epithelial cells, as was later proposed (Frank et al., 2019)? While the current paper does not directly address the question of bipotential progenitors, it does suggest that a significant proportion (around 20%) of committed (at E18.5) AT1 and AT2 cells derive from progenitors expressing an opposing alveolar epithelial cell marker (Fig.8A). This calls into doubt the model of early alveolar lineage specification, which posits that the vast majority of alveolar cells are lineage committed as early as E13.5 (Frank et al., 2019). Regardless of whether the 20% of E14.5-E15.5 lineage-labeled cells shown to express an alternate lineage marker at E18.5 derived primarily from bipotential progenitors or represent transdifferentiating cells, they do indicate that a significant minority of mature alveolar epithelial cells arise from lineage-flexible progenitors.

**Figure 8:**
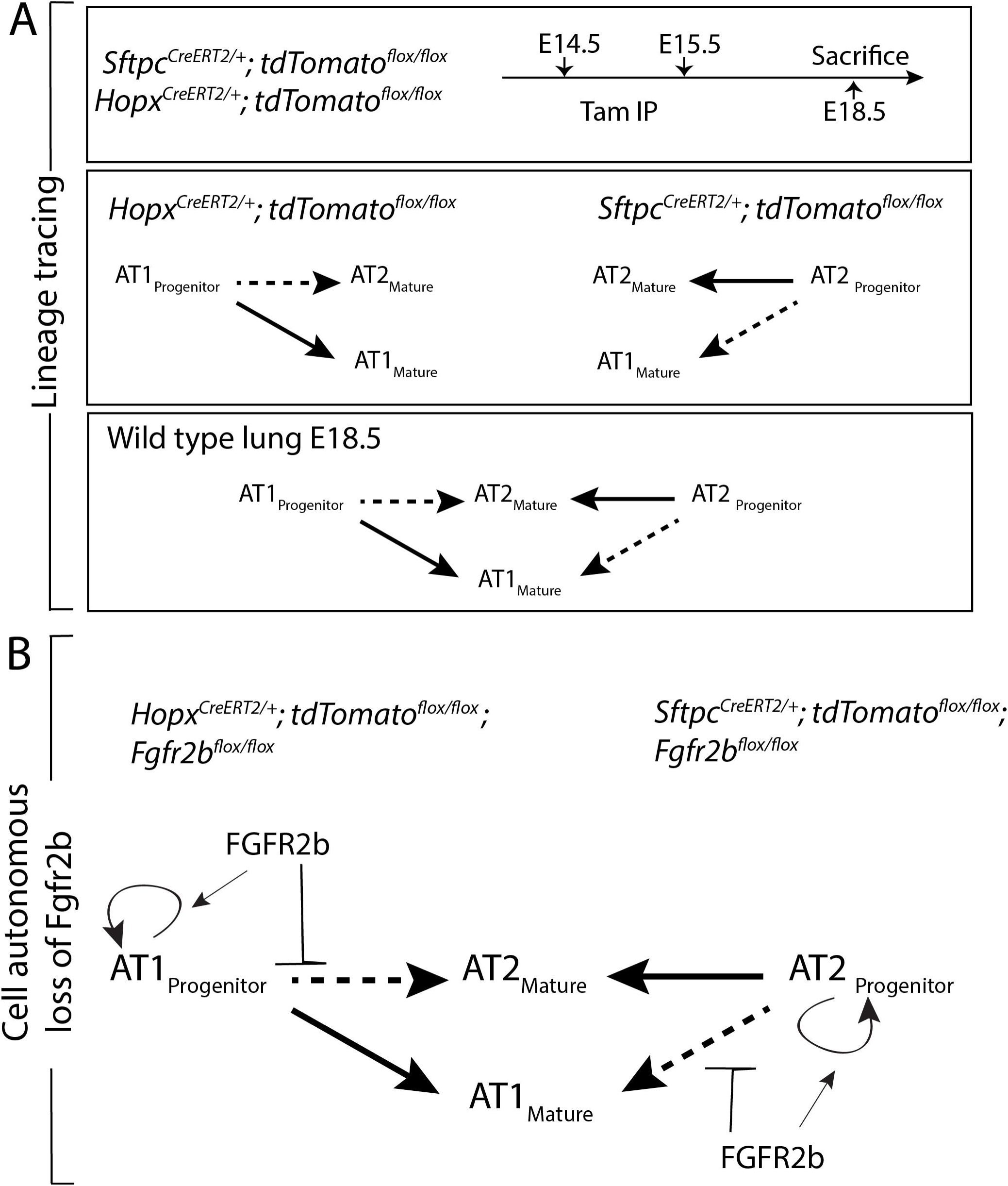
Summary and model of Fgfr2b regulation of AT lineage formation. **(A)** Lineage tracing experiments at E14.5 and E15.5 (late pseudoglandular/early canalicular) embryonic development (first panel) showed that labeled AT1 progenitors give rise primarily to mature AT1s at E18.5, but do contribute a significant minority to the AT2 lineage, while the reverse is true for labeled AT2 progenitors (second panel). Taking the lineage tracing data together, we propose the model in the third panel of AT progenitor contribution to mature AT cells in wild type lungs at E18.5. **(B)** Cell autonomous deletion of Fgfr2b experiments suggest that, apart from the expected impacts of Fgfr2b signalling on AT progenitor proliferation, Fgfr2b *prevents* AT progenitors from actively transdifferentiating to the opposing lineage.

### Role of Fgfr2b signalling on alveolar epithelial lineage formation and maintenance

Little is known concerning the role of Fgfr2b signalling on the development and maintenance of alveolar cells. Most of the research on this topic has looked at AT2 cells in the context of adult homeostasis and repair after injury. For example, using the bleomycin-induced lung fibrosis model, it was reported that mice with specific deletion of Fgfr2 in SFTPC-positive AT2 cells were less able to repair after injury, showed increased mortality, and had fewer AT2 cells overall (Dorry et al., 2020). This study also showed that even in the absence of injury, Fgfr2 deletion resulted in enlarged airspaces and increased collagen deposition, as well as a decrease in the number of AT2 cells, suggesting that Fgfr2 is required for AT2 maintenance.

These results support earlier work which looked at the loss of *Etv5* in AT2 cells during homeostasis and repair after bleomycin-induced lung injury (Zhang et al., 2017). Here, it was shown that Etv5 is required to maintain AT2 cells, for in its absence, AT2 cells transdifferentiated to AT1s. Furthermore, loss of *Etv5* in AT2 cells drastically impaired the repair process of the epithelium after lung injury, resulting in fewer AT2 cells altogether. It is well established that Etv5 is regulated by Fgfr2b signalling (Herriges et al., 2015), and it was suggested that Etv5 protein stability in AT2 cells is controlled by Ras-mediated Erk signalling (Zhang et al., 2017).

Interestingly, we recently described that miR-142 is critical for the formation of the alveolar lineage and more particularly for the formation of AT1 cells (Shrestha et al., 2019). In the absence of *miR-142*, alveolar progenitors differentiate towards the AT2 lineage in a Ep300 dependent manner while overexpression of *miR-142* in alveolar progenitors leads to their differentiation towards the AT1 lineage. To date, the link between Fgfr2b signaling and *miR-142* is still unclear.

The differentiation of alveolar cells depends not only on ligand-receptor interactions, but also on mechanical forces. For example, it was demonstrated that AT1 cell differentiation is dependent on the mechanical stretching of AT1 progenitors from fetal breathing movements (Li et al., 2018). In the same study, it was shown that some progenitors avoid an AT1 fate due to an Fgf10/Fgfr2b mediated build-up of myosin in the apical region of a proportion of the cells. This increase in myosin is related to the protrusion of these progenitors from the monolayer epithelium, thereby sparing them from the mechanical forces experienced by the cells lining the lumen. These protruded cells adopt an AT2 fate.

Very little is known about the regulation of AT lineage formation by Fgfr2b signalling during development. However, what is emerging supports the evidence seen during homeostasis and repair after injury. That is, that Fgfr2b signalling maintains AT2 cell identity. It seems clear that even during early pseudoglandular development, Fgfr2b signalling primarily affects AT2 lineage formation. As early as E12.5, for example, our lab had observed that just after nine hours Fgfr2b ligand inhibition the AT2 signature found in distal tip progenitors was significantly downregulated (Jones et al., 2019). However, this is only part of the story. The data presented in the current paper strongly suggests that Fgfr2b signalling not only affects the AT2 lineage, but the AT1 lineage as well. The cell autonomous loss of Fgfr2b in labeled SFTPC^Pos^ AT2 progenitors or in HOPX^Pos^ AT1 progenitors leads to an *increase* in the ratio of labeled cells belonging to the alternate lineage (Figs. 4, 5). While it is true, since Fgfr2b signalling also has a proliferative effect at the timepoint of labelling (E14.5-E15.5, (Jones et al., 2020)), that the absolute number of labeled AT2 or AT1 cells in our Fgfr2b knock-out lineage tracing experiments decreased, the relative ratios of labeled AT2 or AT1 progenitors producing labeled AT1 or AT2 progeny, respectively, increased. These data suggest that Fgfr2b signalling, while acting primarily on the AT2 lineage at later stages of development, serves to maintain the lineage commitment of both AT2 and AT1 progenitors during late pseudoglandular and early saccular lung development (see model in Fig.8B).

### Narrowing of Fgfr2b signalling to an AT2 sub-population over embryonic development highlights the heterogeneity of AT2 cells

Assessing, at a fixed stage of development (e.g. E17.5), the expression patterns of the Fgfr2b signatures from earlier embryonic stages revealed that the closer these signatures came to the fixed timepoint, the more narrowly confined their expression became. The earlier in embryonic development one looks, the less differentiated the Fgfr2b-responding progenitors will be. Thus, at E12.5, for example, the distal tip epithelial progenitors are largely uncommitted multipotent cells; as a consequence, Fgfr2b signalling at earlier stages affects the differentiation of an increasing number of cell lineages. However, as development progresses, distal progenitors become more lineage-restricted, not only as a consequence of ligand-receptor interactions and tissue geometry, but also in response to mechanical and physical forces.

Our data indicate that as alveolar cell lineages emerge and develop (beginning as early as E12.5), the role of Fgfr2b signalling shifts, eventually concentrating in a sub-population of the AT2 lineage (Fig. 6). This highlights an increasingly appreciated fact that cellular populations are heterogeneous, and points to the need to identify and further classify these sub-populations of cells (Chen and Liu, 2020). Indeed, recent work from our lab has identified two major sub-populations of AT2 cells in adult mice: those expressing high levels of Sftpc and Fgfr2b, and those expressing low levels of both markers in addition to high expression of a cell surface protein called PD-LI (Ahmadvand et al. 2021). It was shown that this latter immature sub-population of AT2s is quiescent during homeostasis, but becomes activated during the repair response to pneumonectomy, eventually expanding to replenish the mature AT2 population. The engagement of this quiescent population in response to injury likely depends on Fgfr2b signalling. In line with this idea, the evidence presented in the current study suggests that AT2 cells can be sub-classified according to responsiveness to Fgfr2b signalling. Whether this responsiveness is a consequence of geographic proximity to zones of active Fgfr2b ligand activity, or whether a subset of cells are no longer capable at a molecular level to respond to Fgfr2b ligands, is yet to be determined. What is evident from the current study, however, is that a subset of AT2 cells, which is activated after injury to replenish the AT1 population, highly expresses E16.5 Fgfr2b signature genes (Fig. 7). By mechanisms still unknown, as these activated AT2s transition to AT1s through a Krt8-positive airway differentiation intermediate state, the Fgfr2b signature is effectively shut off. Thus, the responsiveness of alveolar cells to signalling pathways, in this case Fgfr2b, does demarcate populations of cells critical for repair after injury, and likely during development, that should be considered in more refined classification schemes.

## Supplementary figures

**Figure S1:**
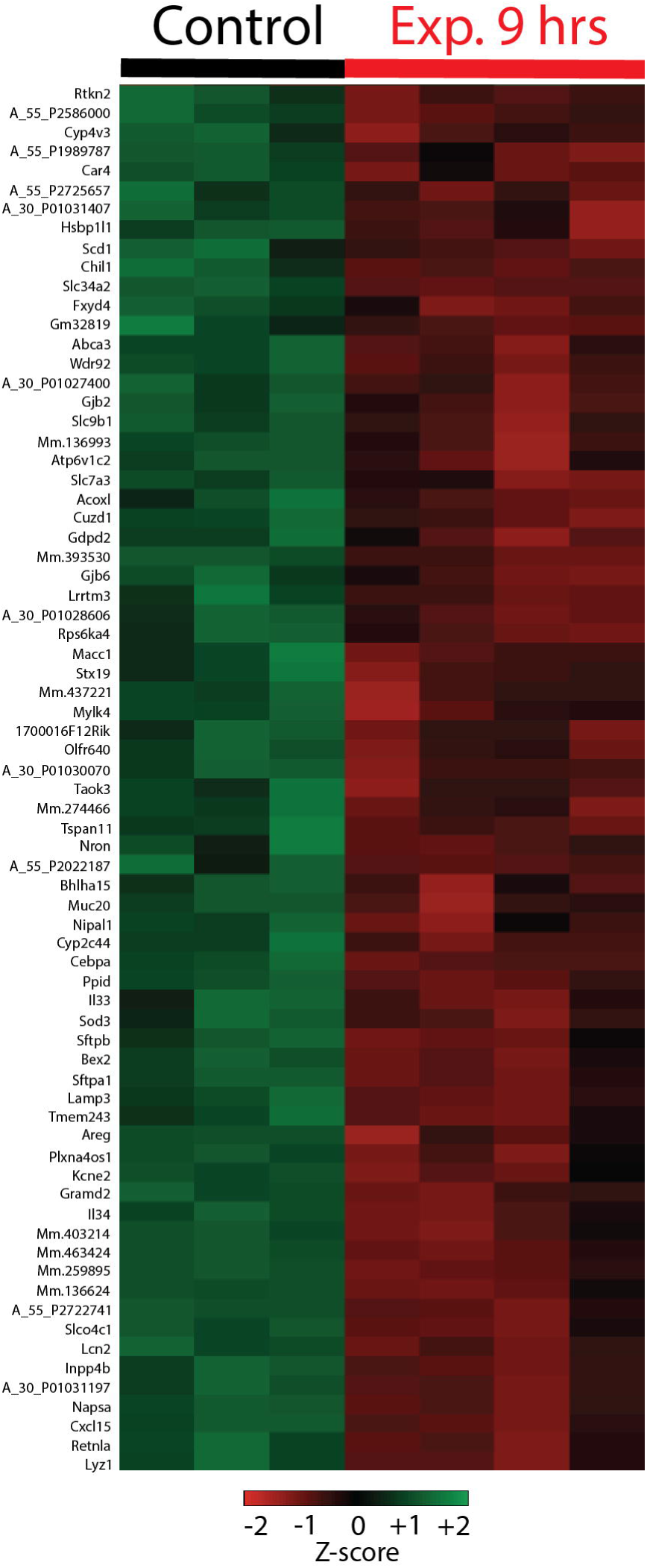
Fgfr2b signature genes at E16.5. **(A)** Heatmap, from Figure 1, showing the 72 downregulated genes after 9 hrs. Fgfr2b ligand inhibition. These genes comprise the Fgfr2b signature at this time-point. **(B)** *In situ* hybridization expression patterns from E14.5 embryonic sections retrieved from the GenePaint database (https://gp3.mpg.de/). Only a small number of the 72 regulated genes were found in the database, and of those a limited number showed clear expression patterns.

**Figure S2:**
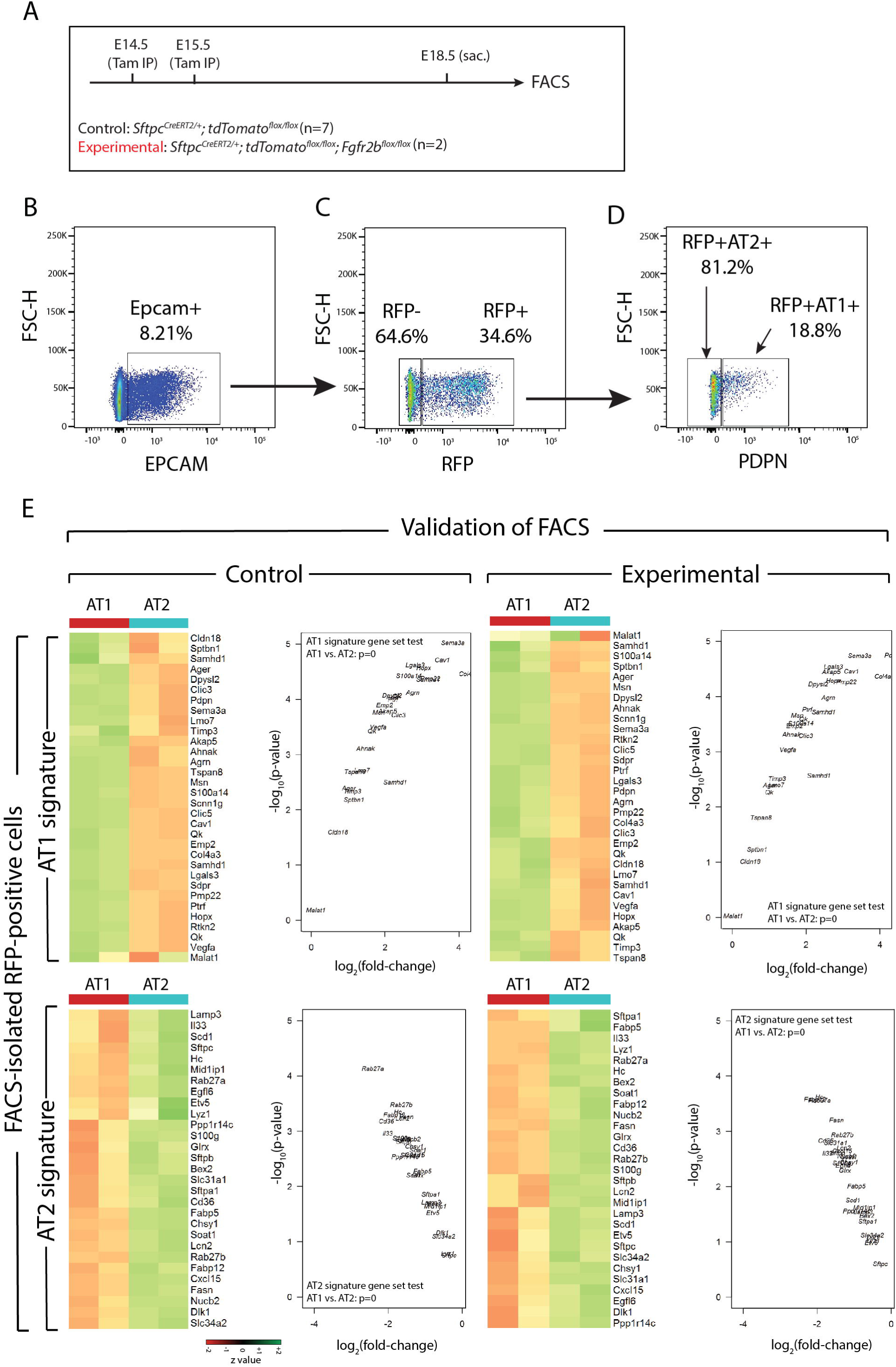
Quality control analysis of FACS-isolated RFP-labeled AT1 and AT2 cells. **(A)** Heatmaps and volcano plots showing AT1 and AT2 signature gene expressions in FACS-isolated RFP-labeled AT1 and AT2 cells from control and **(B)** experimental lungs. Gene-set tests confirm that isolated AT1 and AT2 cells are highly enriched in their canonical signature genes.

**Figure S3:**
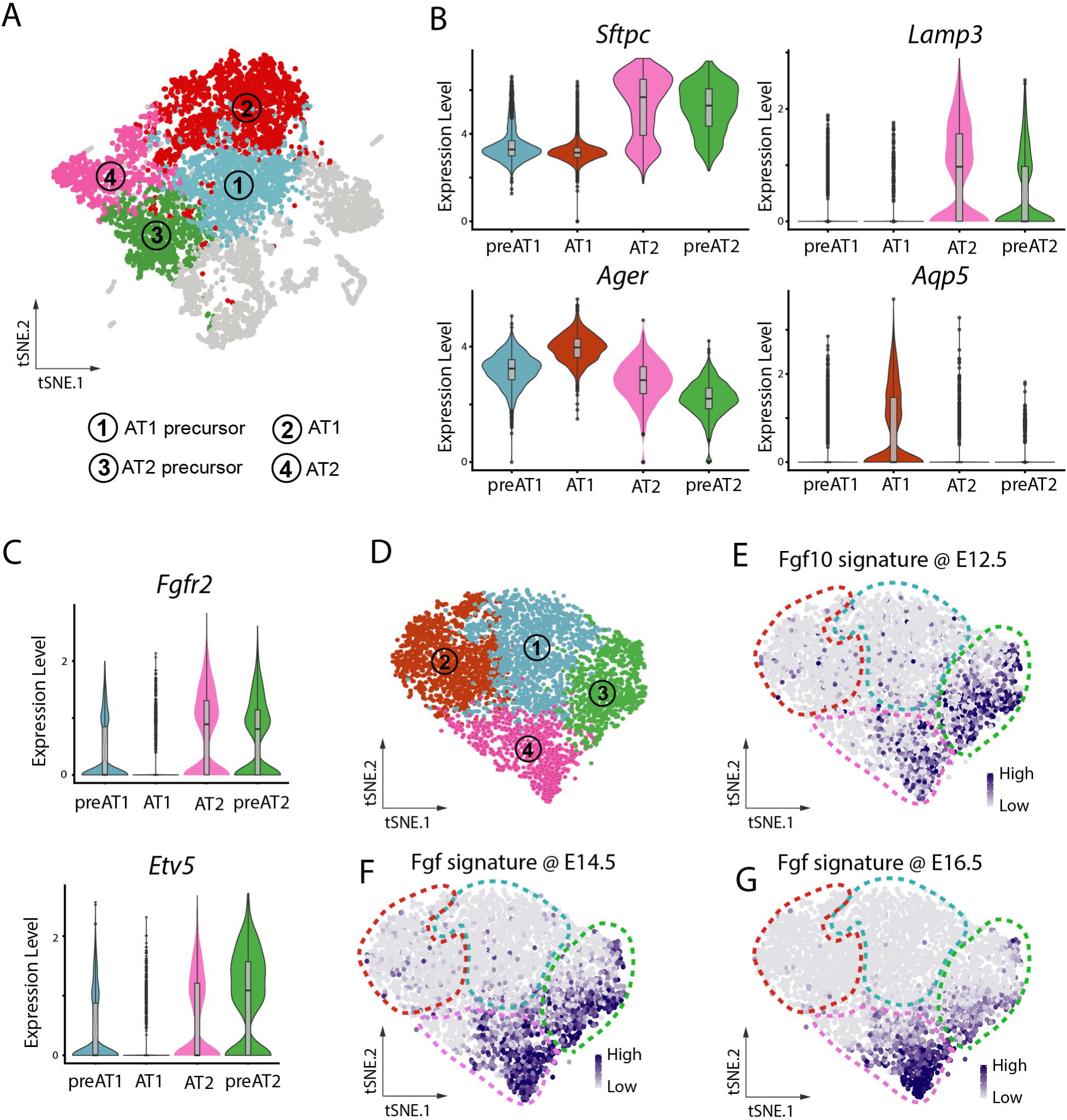
Data-mining of scRNA-seq data from isolated E17.5 Nkx2-1-positive cells shows a narrowing of embryonic Fgfr2b signatures to a subcluster of AT2 cells. **(A)** Four clusters reproduced from Frank et al., 2019, GEO GSE113320: 1, blue – AT1 precursor; 2, red – mature AT1 cells; 3, green – AT2 precursor; 4, pink – mature AT2 cells. **(B)** Violin plots depict the expressions of AT2 (*Sftpc* and *Lamp3*) and AT1 markers (*Ager* and *Aqp5*) in the four clusters. **(C)** Violin plots of *Fgfr2* and the canonical downstream effector of Fgfr2b signalling, *Etv5*, show that the two AT2 clusters, as well as to a lesser extent the AT1 precursors, show expression of these genes. **(D-G)** Reorientation of the four clusters from ‘A’ for ease of analysis. Expressions of Fgfr2b signature genes at E12.5 (E), E14.5 (F), and E16.5 (G) reveal a narrowing of Fgfr2b responsive genes to a subcluster of mature AT2 cells (cluster 4) and AT2 precursors (cluster 3).

**Figure S4:**
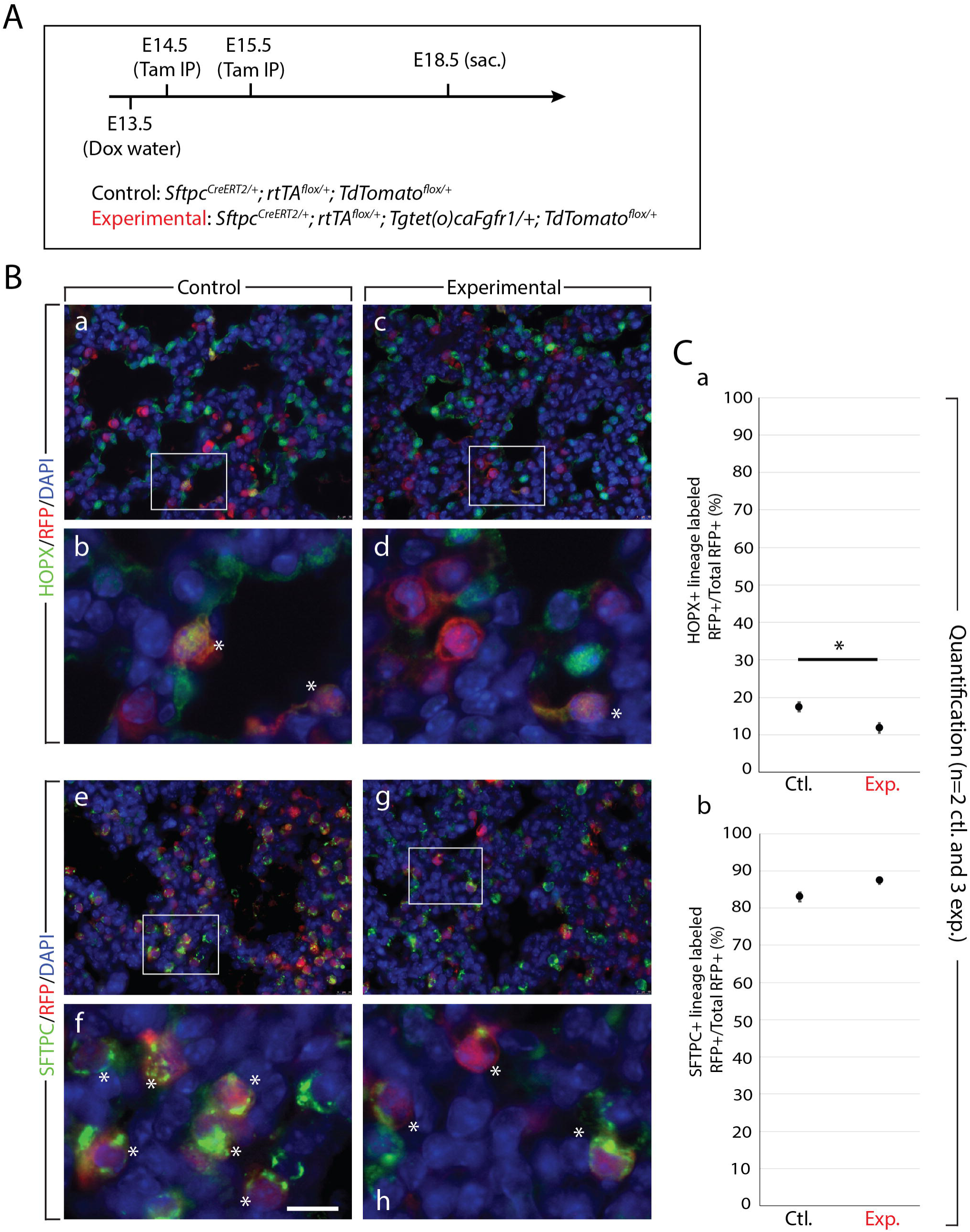
Constitutive expression of Fgfr1 in AT2 progenitors limits transition to AT1 cell fate. **(A)** Experimental design. Timed-pregnant females carrying control (*Sftpc^CreERT2/+^; rtTA^flox/+^; Tomato^flox/+^*) and experimental (*Sftpc^CreERT2/+^; rtTA^flox/+^; Tg^tet(o)caFgfr1/+^; Tomato^flox/+^*) embryos were fed doxycycline water from E13.5 onward. At E14.5 and at E15.5, females were Tam-IP injected. After tamoxifen injection, Cre-based recombination of the floxed *TdTomato* reporter, as well as the floxed *rtTA*, was achieved in AT2 progenitor cells. In experimental embryos, rtTA/doxycycline induces the expression of a constitutively active form of *Fgfr1*, which, when expressed in alveolar epithelial cells, mimics the activity of Fgfr2b signalling. Embryos were sacrificed and lungs harvested at E18.5. **(B)** qPCR. **(C)** Immunofluorescence staining of either AT1 cells (HOPX; panels a-d) or AT2 cells (SFTPC; panels e-h) in control (a, b and e, f) and experimental (c, d and g, h) lineage-labeled (RFP) samples. Asterisks indicate double positive cells. *Scale bar:* 30 µm (a, c, e, g); 75 nm (b, d, f, h). **(D)** Quantification of samples from (C). Graphs show the percentage of double positive cells either from the pool of total lineage-labeled cells (a and c) or from the pool of total AT1 or AT2 cells (b and d). Constitutive expression of Fgfr1 in AT2 progenitors results in a significant decrease from around 18% to 12% in the percentage of labeled AT1 cells over total labeled cells (HOPX^pos^ RFP^pos^/RFP^pos^, graph a), whereas the percentage of labeled AT2 cells trends upward (graph c). Graph b shows little change in the percentage of labeled AT1 cells over total AT1 cells, whereas graph d shows that the percentage of labeled AT2 cells over total AT2 cells is trended upward, suggesting an increased proliferative effect in labeled cells (n=2 control and 3 experimental).

